# Myospreader improves gene editing in skeletal muscle by myonuclear propagation

**DOI:** 10.1101/2023.11.06.565807

**Authors:** Kiril K. Poukalov, M. Carmen Valero, Derek R. Muscato, Leanne M. Adams, Heejae Chun, Young il Lee, Nadja S. Andrade, Zane Zeier, H. Lee Sweeney, Eric T. Wang

## Abstract

Successful CRISPR/Cas9-based gene editing in skeletal muscle is dependent on efficient propagation of Cas9 to all myonuclei in the myofiber. However, nuclear-targeted gene therapy cargos are strongly restricted to their myonuclear domain of origin. By screening nuclear localization signals and nuclear export signals, we identify “Myospreader”, a combination of short peptide sequences that promotes myonuclear propagation. Appending Myospreader to Cas9 enhances protein stability and myonuclear propagation in myoblasts and myofibers. AAV-delivered Myospreader dCas9 better inhibits transcription of toxic RNA in a myotonic dystrophy mouse model. Furthermore, Myospreader Cas9 achieves higher rates of gene editing in CRISPR reporter and Duchenne muscular dystrophy mouse models. Myospreader reveals design principles relevant to all nuclear-targeted gene therapies and highlights the importance of the spatial dimension in therapeutic development.

## Introduction

Gene editing approaches for skeletal muscle have employed numerous Cas9-based strategies, including dual cuts (*1, 2*), indel formation (*3, 4*), base editing (*5*), prime editing (*6, 7*), transcriptional modulation (*8*), and epigenetic modulation (*9*) to restore gene expression or eliminate/repress toxic sequences (*10*). However, proof of concept studies in mouse (*1, 11*) and large animal models (*2, 4*) have required extremely high doses of AAV to achieve therapeutic benefit. The syncytial nature of myofibers likely requires transduction of many myonuclei within the cell to achieve high rates of editing. Novel muscle-tropic capsids have improved transduction efficiency (*12*), but still only a fraction of myonuclei per myofiber are transduced (Fig. S1A and S1C). Critically, the inability of Cas9-based editors to propagate efficiently to non-transduced nuclei has led to surprisingly few myonuclei containing Cas9 protein (*13*) and is partly responsible for the sparse restoration of dystrophin in Dmd-edited myofibers (*14, 15*). While small cytoplasmic proteins diffuse well throughout the myofiber (Fig. S1B), mRNA (*16–18*) and nuclear proteins are more limited in their ability to propagate from their progenitor myonucleus to other neighboring myonuclei, with larger proteins propagating less effectively (*19, 20*). S. aureus and S. pyogenes Cas9 (127 kDA and 162 kDa, respectively) (*21, 22*) are usually tagged with dual nuclear localization sequences (NLSs) to enhance nuclear targeting (*23*). This feature, while important for efficient editing, results in poor myonuclear propagation due to the size of the proteins, limiting overall editing efficiency.

Numerous endogenous transcription factors and RNA binding proteins use a combination of NLS and nuclear export signals (NESs) to facilitate nucleo-cytoplasmic shuttling (*24–26*). Here, we test the hypothesis that bidirectional nuclear-cytoplasmic transport can broaden the myonuclear propagation profile of Cas9, resulting in overall improved editing efficiency. We screened combinations of NLS, NES, and other elements to promote propagation and localization to as many nuclei as possible in a syncytial environment. We identify one combination, “Myospreader”, that promotes Cas9 myonuclear propagation across myofibers, enhances protein stability, and increases muscle gene editing efficiency *in-vivo*. This concept is applicable to any gene therapy cargo designed to target myonuclei in the context of sparse delivery and highlights the importance of considering the spatial dimension in gene regulation, editing, and therapy.

### Combining nuclear import and export signals promotes protein shuttling to a greater number of myonuclei in myotubes and myofibers

Multiple observations suggest that AAV gene therapies encoding nuclear-targeted cargoes are restricted to progenitor myonuclei following mRNA export, translation, and subsequent re-import of the encoded protein (*13–15*). Consistent with this, we observed that a limited number of myonuclei in WT C57BL/6 mice are functionally transduced by AAV packaged with a GFP transgene (Fig. S1A, S1C), and exogenous mRNAs originating from these myonuclei are restricted to their respective myonuclear domains (Fig. 1A). Inspired by nucleocytoplasmic shuttling of endogenous proteins (*24–26*), we hypothesized that appending both a nuclear localization sequence (NLS) and nuclear export sequence (NES) onto cargoes may permit escape from progenitor nuclei and facilitate propagation to distant myonuclei (Fig. 1B). Using GFP as a reporter of protein localization, we screened multiple NLS/NES combinations (Table S1)(*27*) by establishing stable C2C12 myoblasts expressing each reporter and fusing them to naïve C2C12 myoblasts on micropatterned gelatin (*28*). Chimeric myotubes were then imaged to assess GFP signal in myonuclei (Fig. 1C). The balance of NLS and NES strengths influenced the overall localization pattern and propagation of GFP (Fig. 1D, Fig. S1D), where stronger overall export behavior resulted in predominantly cytoplasmic localization, while stronger overall import behavior yielded nuclear localization of GFP but a limited number of GFP-positive nuclei. The best candidate, which showed significantly more nuclear and overall GFP signal than all others (Fig. 1E, Fig. S1E and S1F), encoded an attenuated SV40 NLS on the N-terminus and both an Alyref NES and HIV rev nucleolar localization signal (NoLS) on the C-terminus; we termed this combination “Myospreader”.

**Figure 1.**
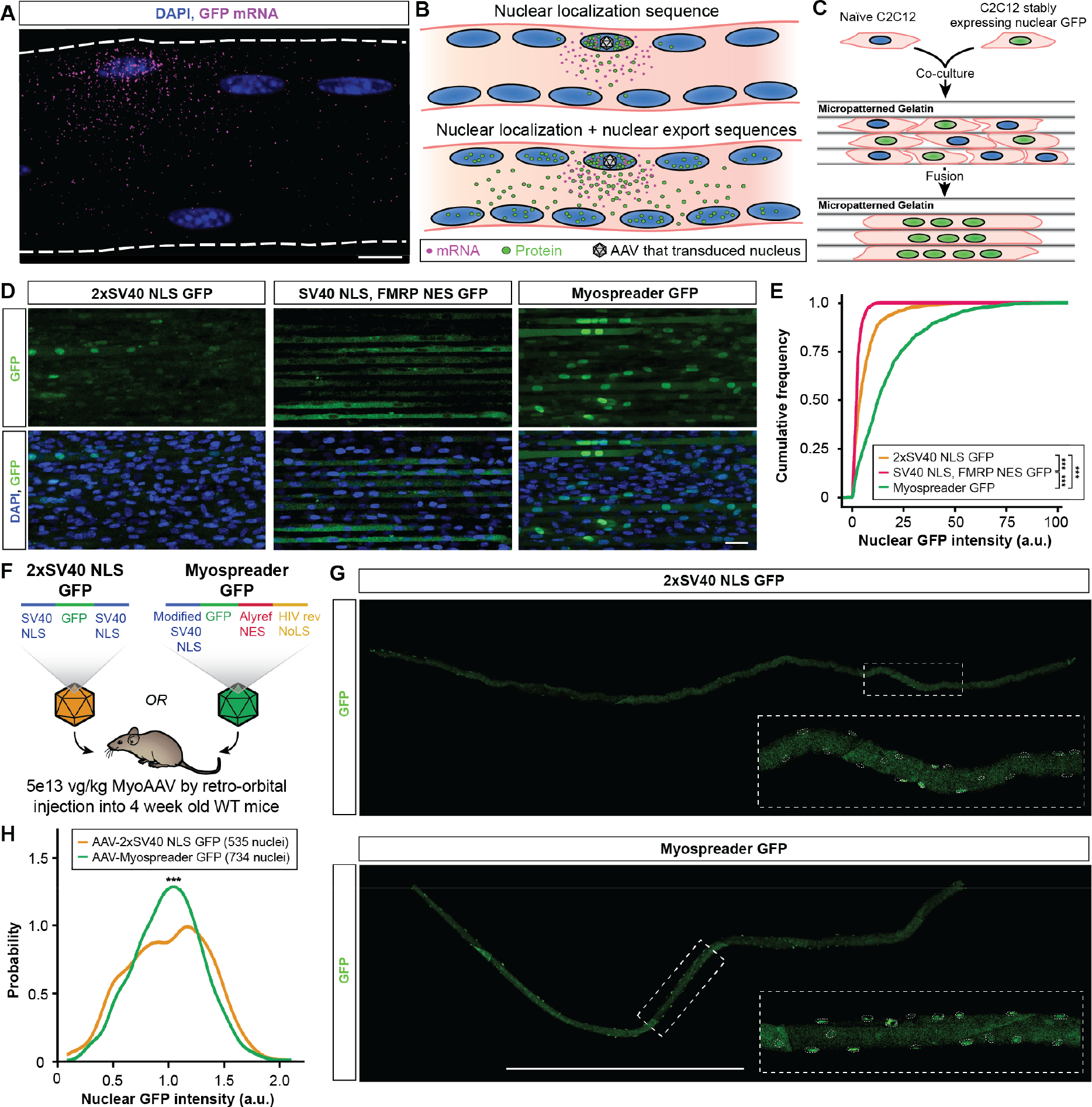
Nuclear import and export sequence combinations enhance myonuclear GFP trafficking in myotubes and myofibers. (**A**) HCR-FISH against GFP mRNA (magenta) in a tibialis anterior myofiber of a mouse treated with an AAV encoding GFP. Scale bar: 10 µm. (**B**) Model for why protein cargoes with nuclear localization sequences accumulate in certain nuclei and not others. Peptide tags that facilitate both nuclear import and export may improve trafficking of protein cargoes to non-progenitor nuclei. (**C**) Schematic of C2C12 fusion experiment to identify NLS/NES combinations that improve propagation of GFP across multiple myonuclei. (**D**) Representative images of GFP fluorescence (green) in chimeric C2C12 myotubes expressing NLS/NES combinations. Scale bar: 40 µm. (**E**) Cumulative distribution functions of nuclear GFP signal for NLS/NES combinations in chimeric C2C12 myotubes. Statistical significance by Kolmogorov-Smirnov test. (**F**) Schematic of experiment to assess nuclear propagation of GFP in 4-week-old WT mice treated systemically with 5E+13vg/kg 2xSV40 NLS GFP AAV or Myospreader GFP AAV. (**G**) GFP signal (green) in representative TA myofibers isolated from treated mice. Myonuclei borders indicated with dashed lines. Scale bar: 1 mm. (**H**) Density plot of nuclear GFP signal in TA myofibers of treated mice. Statistical significance by Kolmogorov-Smirnov test.

To assess the performance of Myospreader *in vivo*, we packaged CBH promoter-driven 2xSV40 NLS GFP or Myospreader GFP using MyoAAV (*12*) and systemically administered 5e13 vg/kg virus to 4 week old C57BL/6 mice. After 4 weeks, tibialis anterior (TA) myofibers were isolated and imaged (Fig. 1F). Myospreader GFP displayed much more uniform localization of GFP protein across myonuclei of myofibers, in contrast to the patchy, infrequent nuclear GFP signal with the 2xSV40 NLS GFP construct (Fig. 1G). Indeed, the distribution GFP intensity of myonuclei, normalized by average GFP signal per myofiber, showed a more uniform distribution across myonuclei as compared to 2xSV40 NLS GFP (Fig. 1H).

### Adding Myospreader to Cas9 enhances protein stability and localization patterns in myoblasts and myofibers

Next, we appended the Myospreader NLS/NES tags to *S. aureus* Cas9 (SaCas9) and assayed its localization pattern in C2C12 myoblasts. Similar to 2xSV40 NLS Cas9, Myospreader Cas9 maintains nucleolar enrichment, a pattern also observed with other Cas9 variants (*29*). However, Myospreader Cas9 also showed some cytoplasmic localization (Fig. 2A). To confirm that Myospreader does not reduce Cas9 editing efficiency or stability in mononucleated cells, we transiently transfected both constructs into a C2C12 myoblast line that stably expresses the Ai14 reporter (*30*) and gRNAs that target the 3 polyadenylation sites upstream of tdTomato; successful excision of this region with a double cut and non-homologous end joining (NHEJ) results in increased tdTomato mRNA and protein expression (Fig. 2B). Both 2xSV40 NLS Cas9 and Myospreader Cas9 showed similar tdTomato and Cas9 mRNA expression levels as assessed by RT-qPCR (Figs. 2C and D), but Myospreader Cas9 showed greater relative levels of Cas9 protein, as assessed by Western blot. This suggests that Myospreader tags also enhance stability of the protein (Fig. 2E).

**Figure 2.**
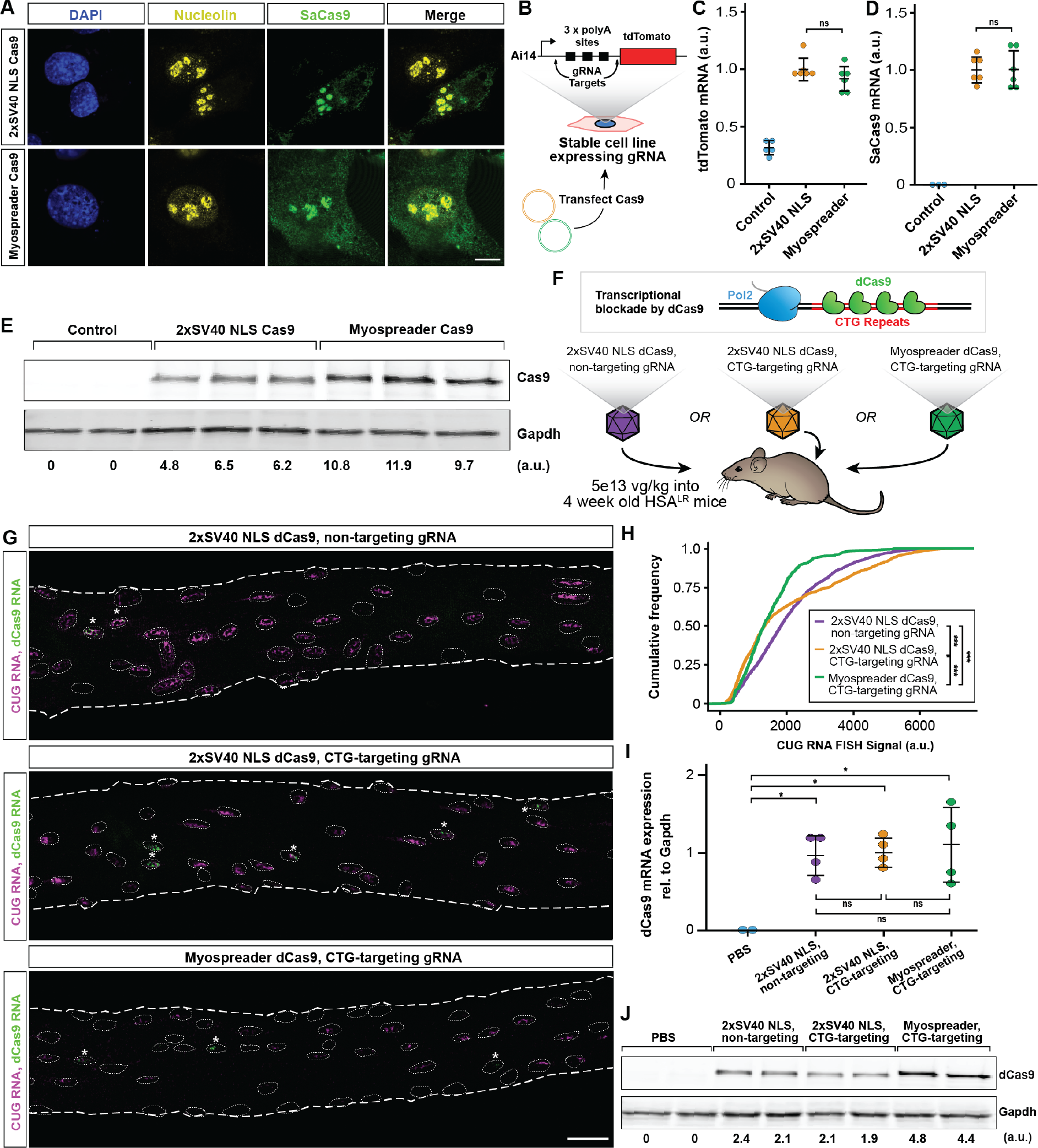
Myospreader improves stability and localization of Cas9/dCas9 in myoblasts and myofibers. (**A**)Representative IF images of C2C12 myoblasts transfected with plasmids encoding 2xSV40 Cas9 or Myospreader Cas9. SaCas9 is shown in green, nucleolin in yellow, DAPI in blue. Scale bars: 10 µm. (**B**) Schematic of experiment to assess Myospreader Cas9 stability and editing efficiency in C2C12 myoblasts. (**C**) tdTomato mRNA expression as assessed by RT-qPCR in AI14 reporter C2C12s following transfection with plasmids encoding 2xSV40 Cas9 or Myospreader Cas9. Plotted relative to mean of the 2xSV40 NLS Cas9 group. Significance by Student’s t-test. (**D**) Cas9 mRNA expression as assessed by RT-qPCR in AI14 reporter C2C12s following transfection with plasmids encoding 2xSV40 Cas9 or Myospreader Cas9. Plotted relative to mean of the 2xSV40 NLS Cas9 group. Significance by Student’s t-test (**E**) Western blot against Cas9 and GAPDH in Ai14 reporter C2C12s following transfection with plasmids encoding 2xSV40 Cas9 or Myospreader Cas9. Relative signal intensity determined by densitometry at the bottom. A.U.: Arbitrary Unit, normalized to GAPDH. (**F**) Schematic of dCas9 impeding transcription of toxic CUG RNA and experiment to measure efficacy of 2xSV40 NLS dCas9 with non-targeting gRNA AAV, 2xSV40 NLS dCas9 with CTG-targeting gRNA AAV, or Myospreader dCas9 with CTG-targeting gRNA. Each treatment was packaged in a single AAV vector systemically delivered to 4-week-old HSA^LR^ mice at a dose of 5E+13 vg/kg. (**G**) HCR-FISH to detect CUG RNA foci (magenta) and dCas9 mRNA (green) in myonuclei of representative TA myofibers isolated from treated HSA^LR^ mice. Myonuclei borders are indicated with dashed lines. Asterisks indicate dCas9-positive myonuclei. Scale bar: 30µm. (**H**) Cumulative distribution functions of HCR-FISH CUG RNA signal in myonuclei proximal to dCas9-expressing myonuclei in myofibers isolated from treated HSA^LR^ mice. Significance by Kolmogorov-Smirnov test. (**I**) dCas9 mRNA expression as assessed by RT-qPCR in Gastrocnemius muscle of treated HSA^LR^ mice. Plotted relative to mean of the 2xSV40 NLS Cas9 non-targeting group. Significance by Tukey’s HSD. (**J**) Western blot detecting dCas9 and GAPDH in gastrocnemius muscle of treated mice. Relative signal intensity determined by densitometry at the bottom. A.U.: Arbitrary Unit, normalized to GAPDH.

We next sought to investigate how Myospreader might impact the function of Cas9 in muscle *in vivo*. We first chose to test the binding functions of deactivated Cas9 (dCas9) in the HSA^LR^ model of myotonic dystrophy, a setting in which we have previously demonstrated the ability of dCas9 to impede transcription of expanded CTG repeats (*13*). We packaged 2xSV40 NLS dSaCas9 with a non-targeting gRNA, 2xSV40 NLS dSaCas9 with a CTG-targeting gRNA, or Myospreader dSaCas9 with a CTG-targeting gRNA into MyoAAV, and systemically delivered these constructs (5e13 vg/kg) or PBS to 4-week-old HSA^LR^ mice. After 4 more weeks, whole muscle and myofibers were harvested (Fig. 2F). HCR-RNA FISH against dCas9 mRNA sequence and CUG repeats revealed which myonuclei were expressing AAV-derived dCas9 and expanded CUG RNA, respectively. As expected, myofibers from mice treated with the 2xSV40 NLS dCas9 with non-targeting gRNA showed no reduction in CUG RNA in myonuclei expressing dCas9 mRNA or in neighboring myonuclei. Myofibers from mice treated with 2xSV40 NLS dCas9 with CTG-targeting gRNA showed reduction of CUG RNA, but the effect was mostly limited to myonuclei expressing dCas9 mRNA. In contrast, Myospreader dCas9 with CTG-targeting gRNA showed reductions in CUG RNA not only in myonuclei expressing dCas9 mRNA, but also in more distal myonuclei (Fig. 2G, Fig. S2A). Quantification of total CUG RNA signal in myonuclei expressing dCas9 mRNA and their neighbors showed a significant reduction of CUG RNA in the Myospreader dCas9 group compared to the CTG-targeting 2xSV40 NLS dCas9 group (Fig. 2H, Fig. S2A). There was no significant difference in relative dCas9 mRNA expression by RT-qPCR between any of the dCas9 treatment groups (Fig. 2I). However, Myospreader dCas9 showed greater relative protein levels as measured by Western blot than either of the 2xSV40 NLS dCas9 groups, again suggesting that Myospreader increases the stability of dCas9 protein in skeletal muscle (Fig. 2J).

### Myospreader improves Cas9 myonuclear propagation and editing efficiency in-vivo

To investigate whether Myospreader improves gene editing performance by Cas9, we turned to Ai14 reporter mice, in which the tdTomato reporter described above has been integrated into the genome (*30*). We packaged 2xSV40 NLS SaCas9, Myospreader SaCas9, and two Ai14-targeted gRNAs into MyoAAV, and delivered each Cas9-encoding virus paired with the gRNA virus (5e13 vg/kg for SaCas9 virus and 5e13 vg/kg for gRNA virus) systemically into 4 week-old Ai14 mice. Another 4 weeks later, whole muscle and myofibers were isolated (Fig. 3A). Quantification of relative tdTomato mRNA expression by RT-qPCR also showed approximately 2-fold higher levels with Myospreader as compared to 2xSV40 NLS, despite similar overall levels of Cas9 mRNA (Fig. 3B and C). Muscle cryosections from mice treated with Myospreader Cas9 had greater tdTomato signal (Fig. 3D). Levels of tdTomato protein were approximately 2-fold higher in muscles of Myospreader treated mice compared to mice treated with 2xSV40 NLS Cas9 as assessed by western blot (Fig. 3E and Fig. S3C). Levels of Myospreader Cas9 protein measured by western blot were 2-3 fold higher than 2xSV40 NLS Cas9 (Fig. 3F, and Fig. S3D), supporting the theory that spreading Cas9 throughout the sarcoplasm and away from the progenitor nucleus may extend protein half-life. HCR-FISH-IF in myofibers isolated from TA showed improved myonuclear propagation of Myospreader Cas9 protein as compared to 2xSV40 NLS Cas9, and increased abundance in the sarcoplasm. TA myofibers from mice treated with Myospreader Cas9 showed tdTomato mRNA spots in and around multiple adjacent myonuclei, suggesting multiple successful editing events originating from a limited number of Cas9-expressing myonuclei (Fig. S3A). Quantification of tdTomato-positive myonuclei from isolated myofibers showed that Myospreader Cas9 generated significantly more tdTomato mRNA-positive myonuclei relative to the 2xSV40 NLS Cas9 construct (Fig. S3B).

**Figure 3.**
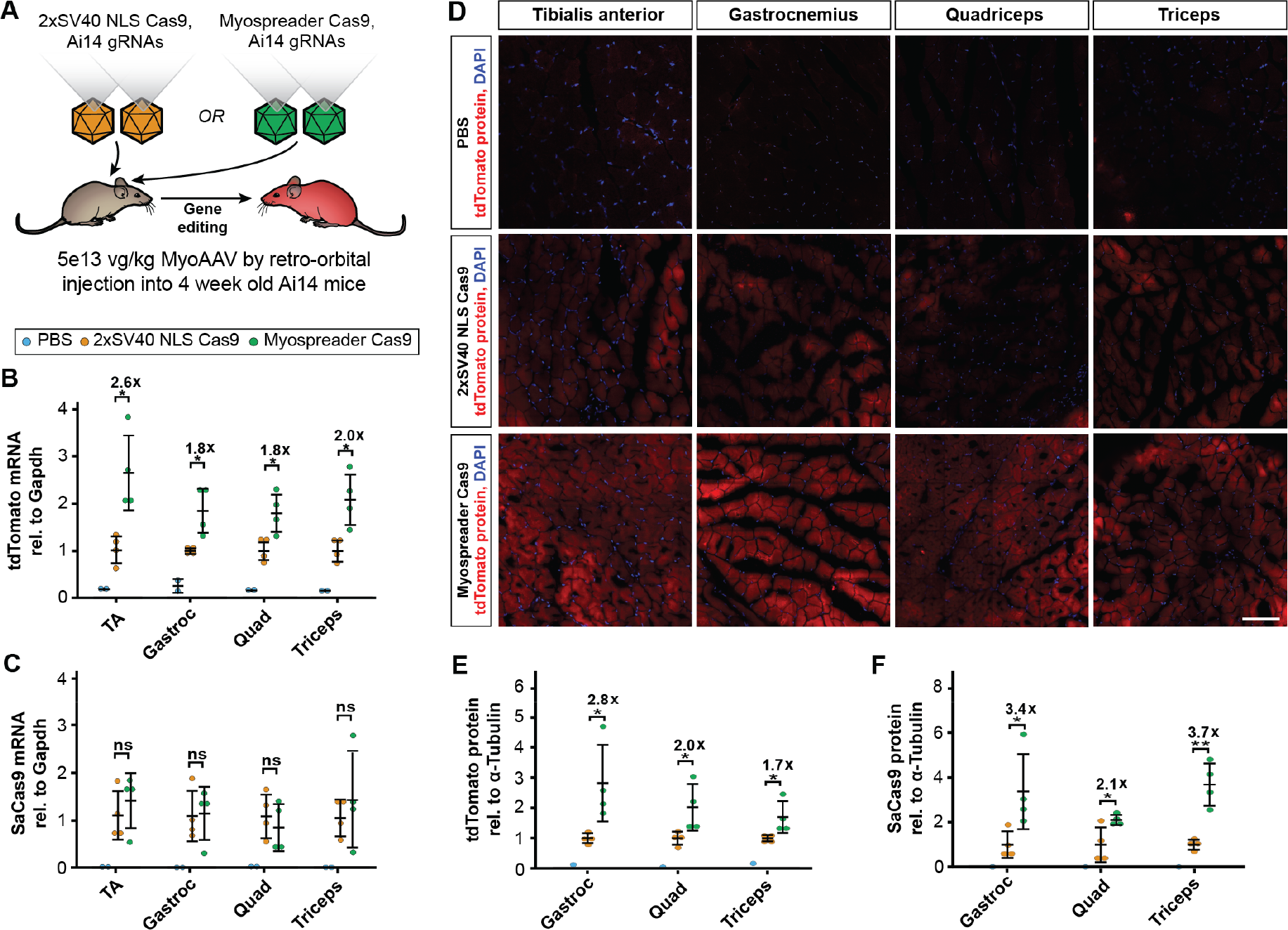
Myospreader improves Cas9 editing efficiency *in-vivo*. (**A**) Schematic of experiment to measure efficacy of AAVs containing 2xSV40 NLS Cas9 or Myospreader Cas9, systemically delivered to 4 week-old Ai14 mice at a dose of 5E+13 vg/kg. (**B**) tdTomato mRNA expression as assessed by RT-qPCR in muscles of treated Ai14 mice. Plotted relative to the mean of the 2xSV40 NLS Cas9 group. Significance by Student’s t-test. (**C**) Cas9 mRNA expression as assessed by RT-qPCR in muscles of treated Ai14 mice. Plotted relative to the mean of the 2xSV40 NLS Cas9 group. Significance by Student’s t-test. (**D**) Representative images of tdTomato fluorescence (red) and DAPI (blue) in muscles of treated Ai14 mice. Scale bar: 100 µm. (**E**) Quantification of Tdtomato protein as determined by Western blot/densitometry, normalized to α-Tubulin. Plotted relative to the mean of the 2xSV40 NLS Cas9 group. Significance by Student’s t-test. (**F**) Quantification of Cas9 protein as determined by Western blot/densitometry, normalized to α-Tubulin. Plotted relative to the mean of the 2xSV40 NLS Cas9 group. Significance by Student’s t-test.

### Myospreader Cas9 increases myonuclear propagation and editing efficiency in a mouse model of Duchenne Muscular Dystrophy

We further assessed the performance of Myospreader Cas9 in the mdx mouse model of Duchenne muscular dystrophy, which contains a nonsense mutation in exon 23 of the *Dmd* gene and expresses little to no dystrophin protein (*31, 32*). Excision of exon 23 with gRNAs targeting the flanking introns can restore reading frame and dystrophin expression (*1, 11*). We packaged 2xSV40 NLS SaCas9, Myospreader SaCas9, and two *Dmd*-targeted gRNAs into MyoAAV, and delivered each Cas9-encoding virus paired with the gRNA virus (5e13 vg/kg for SaCas9 virus and 5e13 vg/kg for gRNA virus) by tail vein into 8 week-old mdx mice. After 4 weeks, whole muscle and myofibers were isolated (Fig. 4A). Myospreader Cas9 treatment produced significantly more exon-23 deleted mRNA in all muscles, with levels ranging from approximately 1.5 to 2-fold higher than 2xSV40 NLS Cas9 despite showing similar Cas9 mRNA transcript levels (Figs. 4B and C). Increases in exon-23 deleted mRNA were supported by Sanger sequencing results showing significantly greater DNA editing in Myospreader Cas9-treated gastrocnemius muscle as compared to 2xSV40 NLS Cas9 (Fig. S4A and S4B). Treatment with Myospreader Cas9 produced significantly higher levels of dystrophin protein and levels of Mysopreader Cas9 protein were also 2 to 4-fold higher than 2xSV40 NLS Cas9 when assessed by western blot (Fig. 4D and E, Fig. S4C and S4D). Immunofluorescence in TA myofibers showed increased abundance and broader myonuclear propagation of Myospreader Cas9 protein as compared to 2xSV40 NLS Cas9, concomitant with increased abundance and broader localization of dystrophin protein (Fig. 4F). Muscles treated with Myospreader Cas9 showed increased dystrophin expression as assessed by immunofluorescence of dystrophin (Fig. 4G). Finally, TA myofibers isolated from mice treated with Myospreader Cas9 displayed more uniform dystrophin expression along the sarcolemma and a decrease in the frequency of centralized myonuclei (Fig. 4H and I). Quantitation of TA myofiber dystrophin revealed that Myospreader Cas9 restores significantly more dystrophin to the myofiber periphery than 2xSV40 NLS Cas9 (Fig. 4J).

**Figure 4.**
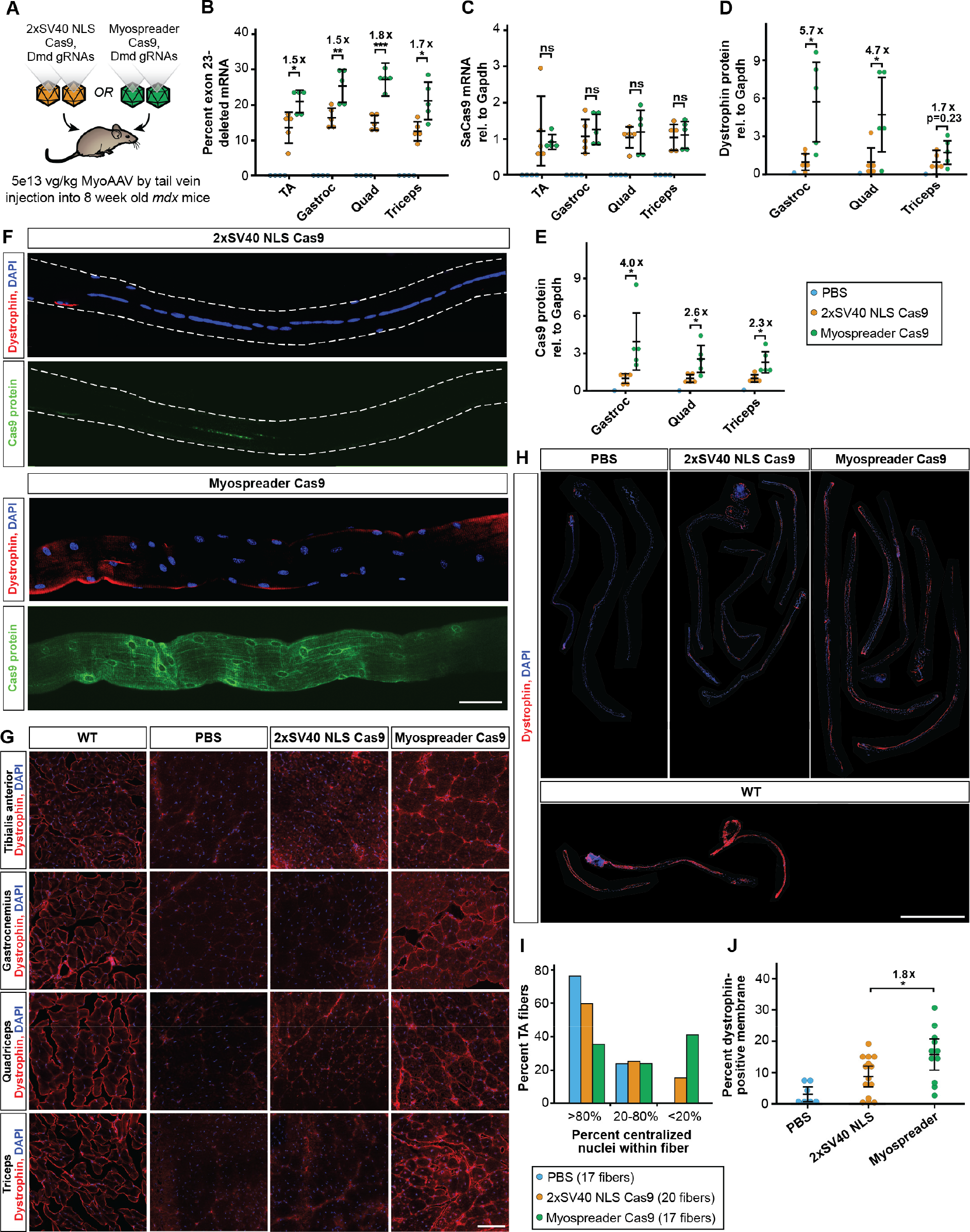
Myospreader improves Cas9 editing efficiency in a Mouse model of Duchenne muscular dystrophy. (**A**) Schematic of experiment to measure efficacy of AAVs containing 2xSV40 NLS Cas9 or Myospreader Cas9, systemically delivered to 4-week-old mdx mice at a dose of 5E+13 vg/kg. (**B**) RT-qPCR quantification of exon-23 deleted *Dmd* mRNA in different muscles of treated mdx mice. Significance by Student’s t-test. (**C**) RT-qPCR quantification of Cas9 mRNA in different muscles of treated mdx mice. Plotted relative to mean of the 2xSV40 NLS Cas9 group. Significance by Student’s t-test. (**D**) Quantification of dystrophin protein as determined by Western blot/densitometry, normalized to GAPDH. Plotted relative to mean of the 2xSV40 NLS Cas9 group. Significance by Student’s t-test. (**E**) Quantification of Cas9 protein as determined by Western blot/densitometry, normalized to GAPDH. Plotted relative to mean of the 2xSV40 NLS Cas9 group. Significance by Student’s t-test. (**F**) IF to detect dystrophin (red) and Cas9 (green) protein in representative TA myofibers isolated from treated mdx mice. Scale bar: 50 µm. (**G**) Representative images of Dystrophin protein (red) and DAPI (blue) in muscle cryosections of treated mdx mice by IF. Scale bar: 100µm. (**H**) IF to detect dystrophin protein (red) and DAPI (blue) in whole TA myofibers isolated from treated mdx mice or untreated 8-week-old WT mice. Scale bar: 1 mm. (**I**) Quantification of centralized nuclei in TA myofibers isolated from treated mdx mice. (**J**) Quantification of dystrophin protein at the periphery of TA myofibers of treated mdx mice or 8-week-old WT mice. Significance determined Student’s T-test.

## Discussion

The syncytial nature of myofibers adds complexity to the conventional view of AAV transduction efficiency and its impact on therapeutic development for muscle. In mononucleated cells, although variation in AAV concatemer number can yield different expression levels, AAV transduction success is binary (*33*). In myofibers, however, successful transduction events occur in a small proportion of myonuclei, leveraging only a fraction of total transgene potential available in the cell. Ultimately, maximum therapeutic benefit will only be achieved if gene therapy cargoes impact all myonuclear domains.

How do size and syncytial nature of myofibers affect AAV transduction? Upon entry into the myofiber, AAV particles may travel along the microtubule network, perhaps visiting several myonuclear domains prior to dynein-dependent recruitment to perinuclear regions and entry into myonuclei (*34, 35*). Similar to endogenous mRNAs, gene therapy mRNAs are transcribed in successfully transduced nuclei, then exported and dispersed throughout the myonuclear domain (*16, 36*). Soluble and structural proteins translated from therapeutic mRNAs can then travel further, the latter perhaps being more restricted (*14, 37*), but nuclear-targeted proteins are trapped close to their progenitor domains (*19, 38*). Large nuclear-targeted proteins then have no way to passively exit the nuclear pore complex and are degraded within the nucleus over time (*39*), similar to other nucleocytoplasmic shuttling proteins (*40*). When combined with low rates of myonuclear transduction, this trapping strongly limits therapeutic efficacy.

By appending an NES, we have facilitated the export of these proteins, extending cytoplasmic residence time and promoting myonuclear propagation. Myospreader, only 29 amino acids longer than 2xSV40 NLS, increases both myonuclear propagation and stability of SaCas9. Indeed, these benefits are probably intertwined; the increased stability of Myospreader Cas9 protein is likely linked to its trafficking behavior. The additional nucleolar localization sequence in Myospreader may help maintain some degree of nucleolar targeting; an intrinsic nucleolar localization signal of SpCas9 was shown to be essential for maintaining protein stability (*29*). Modifications to the specific NLS and NES utilized as well as their N/C-terminus orientation are potential optimization strategies for Myospreader and other NLS/NES combinations. Overall, the combination of cytoplasmic, nuclear, and nucleolar signals in Myospreader SaCas9 may harmonize stability, trafficking, and editing properties; similar principles may have the potential to improve effectiveness of AAV- and mRNA-based therapies in mono- and multi-nucleated cells.

The newest generation of gene editing techniques employ fusions of Cas9 to enzymatic domains to make precise edits to DNA without inducing double strand breaks. Both base and prime editors are large (>197kDa) fusions consisting of a Cas9 fused to DNA targeted enzymatic domains (*41–44*), and additional fusions are being employed for epigenetic editing, transcriptional activation, and transcriptional repression (*8, 9, 45–47*). Many other myonuclear-targeted cargoes are also under development, such as a DUX4 dominant negative for Facioscapulohumeral dystrophy (*48*), Pumilio-PIN fusions for myotonic dystrophy (*49*), and PGC1a for sarcopenia (*50*). All of these approaches can in principle benefit from the enhanced myonuclear propagation and protein stability provided by Myospreader or other NLS/NES combinations. Ultimately, efforts to develop safe and effective gene therapies in any cell type will benefit from considering the spatial dimension of gene regulation.

## Acknowledgements

We thank Sharif Tabebordbar for multiple discussions related to efficiency of gene editing in muscle over the years, and for the eMyoAAV-GFP virus used for experiments in Figure 1A. We thank Ann Kennedey for the mouse illustration used in the figures (doi.org/10.5281/zenodo.3925921). We thank members of the Wang lab for comments and suggestions related to this work.

## Author contributions

Conceptualization: K.K.P., E.T.W. Methodology: K.K.P. Investigation: K.K.P., M.C.V., D.R.M., L.M.A., Y.I.L., H.C. Visualization: K.K.P. Funding Acquisition: E.T.W., H.L.S. Project Administration: K.K.P. Supervision: E.T.W. Resources: E.T.W., Z.Z., H.L.S. Writing – original draft: K.K.P. Writing – review & editing: K.K.P., E.T.W.

## Data and materials availability

All data are available in the manuscript or the supplementary materials. Code is available at https://github.com/kirilpoukalov/Myospreader.git.

## Materials and Methods

### Mice

All animal care and experimental procedures were performed in accordance with the University of Florida Animal Care and Use Committee (IACUC). For the AAV-GFP experiments, 4 week old male and female C57BL/6 mice were injected retro-orbitally with AAV. For the AAV-dCas9 experiment, 4 week old male and female HSA^LR^ mice were injected retro-orbitally with AAV. For the AAV-Cas9-AI14 experiment, 4 week old male and female AI14 mice were injected retro-orbitally with AAV. For the AAV-Cas9-mdx experiment, 8 week old male mdx mice were injected by tail-vein with AAV.

### Cell Lines

The C2C12 mouse myoblast cell line was obtained from ATCC (CRL-1772).

### Constructs

All plasmids were generated using Gibson assembly and In-Fusion cloning according to manufacturer protocols (Takara, 638920). PB-NCT-GFP plasmids were generated using NCT plasmids donated by Zane Zeier and were flanked by PiggyBac transposon arms. AAV-GFP constructs were generated by cloning the CAG promoter, GFP coding sequence, SV40 PolyA signal into a pAAV plasmid backbone. AAV-dCas9 plasmids were generated from the SaCas9 plasmid pX601-AAVCMV∷NLS-SaCas9-NLS-3xHA-bGHpA;U6∷BsaI-sgRNA (Addgene #61591) by incorporation of NLS/NES elements and a CAG-targeting or non-targeting sgRNA. AAV-Cas9 plasmids were generated from SaCas9 plasmid pX601-AAVCMV∷NLS-SaCas9-NLS-3xHA-bGHpA;U6∷BsaI-sgRNA (Addgene #61591) by incorporation of NLS/NES elements and removal of sgRNA and U6 promoter. AAV-Ai14-sgRNA was generated by cloning dual U6 driven sgRNAs targeting flanking regions of Ai14 SV40 polyA tract into a pAAV backbone. AAV-mdx-sgRNA was generated by cloning dual U6 driven sgRNAs targeting flanking introns of *Dmd* exon 23 into a pAAV backbone. Target sequences of gRNAs used are listed in Table S2.

### Stable C2C12 Cell Line Production

A wild type C2C12 myoblast line was co-transfected with PB-NCT-GFP constructs and an mPB plasmid expressing a PiggyBac transposase. Piggybac facilitates stable integration into the host genome(*51*). 48 hours after transfection C2C12s were selected with 5 µg/mL puromycin for 1 week. All transfections were performed using TransIT X2 (Mirus, MIR6003) according to the manufacturer’s instructions.

### Gelatin Hydrogels

8% weight/volume gelatin (Sigma-Aldrich, G1890) and 0.02% chloroform was dissolved in autoclaved water at 65°C. 10% Microbial transglutaminase (mTG) solution was prepared using the heated gelatin solution. mTG gelatin solution was added dropwise onto glass coverslips previously activated using 100 mM NaOH, 0.5% (3-Aminopropyl) trimethoxysilane, and 0.5% glutaraldehyde as described in Bettadapur et al. (*52*). Polydimethylsiloxane (PDMS) stamps were then pressed onto the gelatin and incubated overnight at 37°C. With stamps attached, gelatin was rehydrated with PBS for 1 hour. PDMS stamps were carefully removed from gelatin and the hydrogel was incubated in cell culture media until plating. Cells on hydrogels were fixed with 4% PFA for 10 minutes before imaging.

### Cell Culture

Mouse C2C12 myoblasts were maintained in Dulbecco’s modified Eagle’s medium (DMEM) supplemented with 10% fetal bovine serum and 1% penicillin-streptomycin in a humidified incubator kept at 37 °C and 5% CO2. Once cells reached 70% confluency, they were serum restricted with differentiation media consisting of DMEM supplemented with 2% horse serum and 1% penicillin-streptomycin. Myotubes were imaged approximately 4 days after introduction of differentiation media. Transient transfections were performed using TransIT X2 (Mirus, MIR6003) according to the manufacturer’s instructions. For experiments involving fusion of GFP stably expressing C2C12s with WT C2C12s, cells were plated together at a 1:1 ratio.

### AAV vectors

All AAV plasmids contained AAV2 ITRs. All AAVs were packaged using MyoAAV 2A, with the exception of experiments for Fig. 1F-H, which were packaged using MyoAAV 4A. The vector used in Fig. 1A and Fig. S1A-C was a gift from Sharif Tabebordbar. All other vectors were packaged at SignaGen laboratories and re-titered by qPCR after digestion with Turbonuclease (BPS Bioscience, 50310) for 1 hour at 37 °C and then proteinase K for 2 hours at 50 °C. iTaq Universal Probes Supermix (Bio-Rad, 1725130) was utilized for qPCR reaction. qPCR primers used for titering are listed in Table S3.

### Western Blots

Protein lysates from tissue samples and cells were prepared in HEPES lysis buffer (20mM HEPES, 100mM KCl, pH8.0, 0.1% Igepal, 1x protease inhibitor (Sigma-Aldrich, P8340)). Protein lysates were centrifuged to remove debris. Total protein concentrations were determined using the Pierce 660 protein assay reagent (Thermo Fisher, 22660) and measured on the Nanodrop 2000 (Thermo Fisher). Equal amounts of total protein were loaded per well in each western blot.

Proteins were fractionated in 4–12% Tris-Glycine gel (XP04120BOX, Thermo Fisher). For dystrophin western blots, the first two lanes contain different percentages of wild-type muscle protein diluted in mdx protein from the same muscle to maintain the same total protein as all other lanes. Protein samples were run with NuPAGE™ LDS Sample Buffer (Thermo Fisher, NP0008) and National Diagnostics 10X TRIS-GLYCINE-SDS PAGE Running BUFFER (Fisher, 50-899-90103). All gels were transferred to PVDF membranes using Trans-Blot Turbo RTA Mini LF PVDF Transfer Kits (Bio-Rad, 1704274).

Membranes were blocked for 1 hour in Intercept® Blocking Buffer (LI-COR, 927-60001). All membranes were then incubated for 16 hours at 4°C in primary antibody diluted in blocking buffer. Primary antibodies used were against HA-tag (Cell Signaling Technology 3724S), tdTomato (Takara Bio, 632496), Dystrophin (DSHB, MANEX1011B(1C7), MANEX1011C(4F9), MANHINGE1B clone 10F9), Gapdh (Cell Signaling Technology, 2118L) (abcam, ab83956), and α-tubulin (Sigma-Aldrich, T9026-100UL). For dystrophin western, 3 primary antibodies were used simultaneously. Membranes were washed 3 times in TBST and then incubated with a secondary antibody in blocking buffer for 1 hour at room temperature. Secondary antibodies used were IRDye® 800CW Donkey anti-Mouse IgG (LI-COR, 926-32212), IRDye® 800CW Donkey anti-Rabbit IgG(LI-COR, 926-32213), IRDye® 680RD Donkey anti-Chicken IgG(LI-COR, 926-68075), IRDye® 680RD Donkey anti-Rabbit IgG (LI-COR, 926-68073). Membranes were washed 3 times in TBST after secondary antibody incubation. Membranes were imaged on LI-COR Odyssey imager.

### Cryosectioning

Dissected muscles were flash frozen for 10 seconds in an isopentane bath cooled by liquid nitrogen and stored at -80 °C until sectioning. Cryosectioning was performed on a Leica CM3050 S cryostat microtome. 10 μm sections were adhered to microscope slides and stored at 4 °C until fixation.

### Tibialis Anterior Myofiber Isolation

*Tibialis anterior* (TA) muscle was dissected from 8 or 12 week old mice (4 weeks post injection) of both sexes (only male mdx mice). Muscle was digested with prefiltered 0.02% collagenase in DMEM at 37 °C for 1 h. Digested TAs were agitated by pipetting in prewarmed collagenase DMEM in horse-serum coated plates under microscopy to dissociate individual fibers. Fibers were collected in horse-serum coated plates containing DMEM and stored in 37°C, 5% CO2 tissue culture incubator for no more than 1 hour before fixation.

### HCR RNA smFISH and Immunofluorescence

HCR v3.0 RNA FISH probes, amplifier, and buffers for each target were purchased from molecular instruments. Isolated myofibers, C2C12 myoblasts/myotubes, and muscle sections were processed identically. Samples were fixed in 4% paraformaldehyde (PFA) in phosphate-buffered saline (PBS) for 10 minutes at room temperature, then washed 3 times for 5 minutes with PBS at room temperature. Samples were permeabilized with 1% Triton x-100 for 10 minutes at room temperature. Immunofluorescence was performed before HCR FISH. Samples were blocked in 1% ultra-pure RNase-free BSA (Sigma), 1 U/µL Nxgen RNase inhibitor (Lucigen), and 0.1% Tween 20 (blocking buffer) for 1 hour at RT. Samples were then incubated in primary antibody diluted in blocking buffer for 1 hour at room temperature. Primary antibodies used were against HA-tag (Cell Signaling Technology 3724S, dystrophin (DSHB, MANEX1011B(1C7)), nucleolin (Bethyl Laboratories, A300-711A-T), Aly (Thermo Scientific, MA1-26754). Samples were washed 3 times for 5 minutes with PBS + 0.1% Tween. Samples were incubated with secondary antibodies for 1 hour at room temperature. Secondary antibodies used were Goat Anti-Rabbit IgG H&L Alexa Fluor® 647 (abcam, ab150079), Goat Anti-Rabbit IgG H&L Alexa Fluor® 555 (abcam, ab150086), Goat Anti-Rabbit IgG H&L Alexa Fluor® 488 (abcam, 150081). Samples were washed 3 times for 5 minutes with PBS + 0.1% Tween. If HCR FISH was performed, samples would be fixed with 4% PFA for 10 minutes followed by 3 washes for 5 minutes with PBS and 1 wash for 5 minutes with 2x saline-sodium citrate (SSC) buffer at room temperature. Samples were incubated in pre-warmed HCR hybridization buffer for 30 minutes at 37°C in a humidified chamber. Primary HCR FISH probes were aliquoted into PCR tubes, heated to 95°C for 90 seconds then diluted to 1nM in hybridization buffer and kept at 37°C. Sample hybridization buffer was replaced with hybridization buffer containing probes and samples were incubated overnight in humidified chamber at 37°C. Samples were washed 5 times for 10 minutes with pre-warmer HCR wash buffer and washed 2 times for 5 minutes with 5xSSC + 0.1% Tween (SSCT) and incubated in HCR amplification buffer for 30 minutes at room temperature. During incubation, HCR amplifiers were aliquoted into PCR tubes and incubated at 95°C for 90 seconds, then allowed to cool at room temperature protected from light for 30 minutes. Amplifiers were diluted to 60 nM in HCR amplification buffer. Sample amplification buffer was replaced with amplification buffer containing amplifiers and incubated at room temperature for 3 hours. Samples were washed 5 times for 10 minutes with SSCT and 1 time for 5 minutes with PBS containing 0.1µg/mL DAPI, and then mounted on slides.

### RT-qPCR

RNA was isolated from cell and tissue samples using Tri Reagent (Zymo, R2050-1-200) and Direct-zol RNA MiniPrep kit (Zymo, R2052). cDNA was made using iscript cDNA synthesis kit (Bio-Rad, 1708890). A Taqman probe assay against the exon 4-5 junction was used for quantification of total *Dmd* transcripts. A taqman probe assay against exon 22-24 was used to quantify exon 23-deleted transcripts. Taqman probe assays utilized iTaq Universal Probes Supermix (Bio-Rad, 1725130). Sacas9, tdtomato, and dsacas9 transcripts were quantified using SYBR green PCR assays with SsoAdvanced Universal SYBR Green Supermix (Bio-Rad, 1725270). Mouse GAPDH mRNA was used as the house keeping control for quantification of Sacas9, tdtomato, and dsacas9 expression. Standard curves were generated for all RT-qPCR experiments by amplifying varying amounts of the target sequence in each run. Quantification was done based on the standard curves. RT-qPCR primers are listed in Table S3.

### Sanger Sequencing

gDNA was isolated from mouse gastrocnemius using Quick-DNA Microprep Plus Kit (Zymo D4074). gDNA was amplified using CloneAmp™ HiFi PCR Premix (Takara 639298) and dmd Sanger sequencing primers. Sanger Sequencing was performed by Genewiz (Azenta). Inference of CRISPR Edits (ICE) tool (Synthego) was used to obtain discordance values for each base pair read from the sanger sequencing results(*53*). Discordance values from the first 100bp after the gRNA target site were used to calculated average discordance values.

### Imaging of fixed samples

Myoblasts, myotubes and myofibers were imaged on a Zeiss LSM 880 AxioObserver microscope with Airyscan using a Plan-Apochromat 0.8 NA ×20 Objective (Fig. 4), a Plan-Apochromat 1.3 NA ×40 oil objective (Fig.1,3,4,S1), a Plan-Apochromat 1.4 NA ×63 oil objective (Fig. 2-4), or a Plan-Apochromat 1.4 NA ×100 oil objective (Fig. 2). Airyscan processing was performed on all images using the Zeiss ZEN Black software. HSALR myofibers used in transcription impeding analysis (Fig. 2H) were imaged on an Optical Biosystems Stellarvision instrument.

### Image analysis pipeline

All code is written in R and Python 3. Image analysis was also done in ImageJ. For each image, a maximum intensity z-projection array was generated from 3D Airyscan confocal stacks in CZI format. To detect myonuclei, DAPI channel was thresholded using Li’s or Otsu’s method for automatic threshold selection (54, 55). Total and maximum signal intensity of other channels in myonuclei was measured. For Fig 1H, GFP signal for each myonucleus was normalized to the mean GFP signal of all myonuclei of that fiber. For Fig 2G, only myonuclei that were within the same 470µm × 470µm FOV as dCas9 mRNA positive (identified by dCas9 mRNA channel signal) nuclei were included in the analysis. For Fig S3B, Dystrophin signal was measured on a segmented line along the perimeter of each fiber, then the 10% percentile signal intensity was subtracted from all signal values to account for varying background signal between fibers. Percent Dystrophin positive membrane was calculated based on the percent of signal readings above 60 AU for each fiber.

### Statistical analysis

All statistical analysis was performed in R. All data are presented as mean ± 95% confidence interval. For datasets with only 2 groups compared, an unpaired two-tailed student’s t-test was performed. Equal variance and normality were assumed. For Fig. 1E, Fig 1G, Fig 2G Kolmogorov-Smirnov test was performed. For Fig. 2I Tukey’s HSD test was performed.

## Supplemental Figures

**Table S1.**
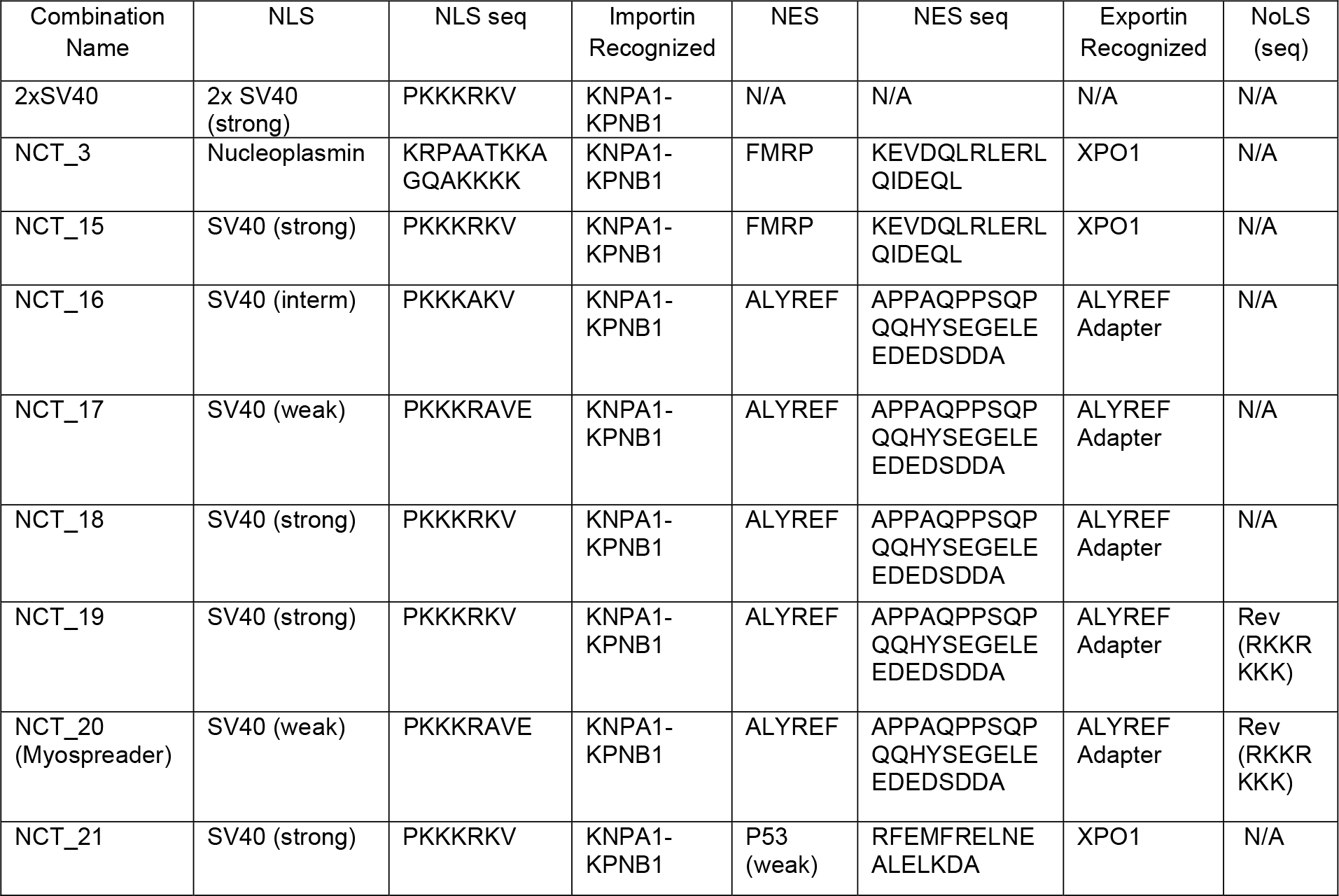
NLS/NES combination tested in mixed nuclei C2C12 myotubes. Related to Figure 1. Table listing all NLS/NES combinations tested in chimeric C2C12 myotubes with the name of each combination, name of each NLS, amino acid sequence for each NLS, importin recognized, name of each NES, amino acid sequence for each NES, exportin recognized, and name of each NoLS.

**Table S2.**
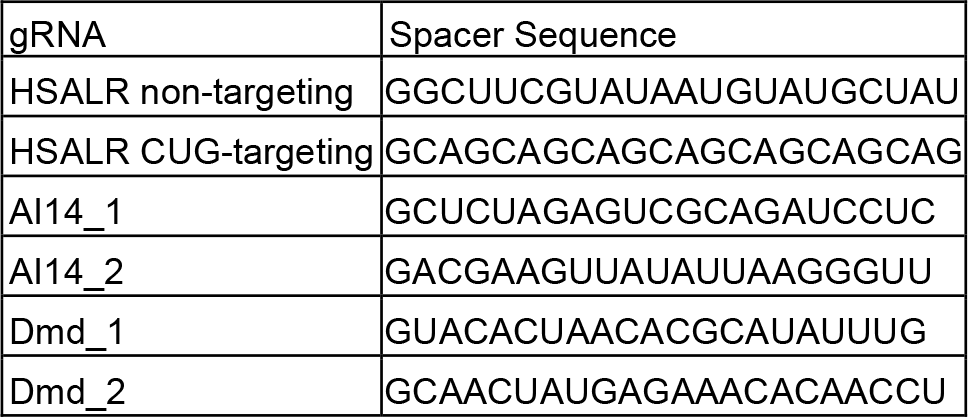
gRNA targets utilized in dCas9 and Cas9 experiments. Table listing names and spacer sequences of all gRNAs used in in-vivo AAV experiments.

**Table S3.**
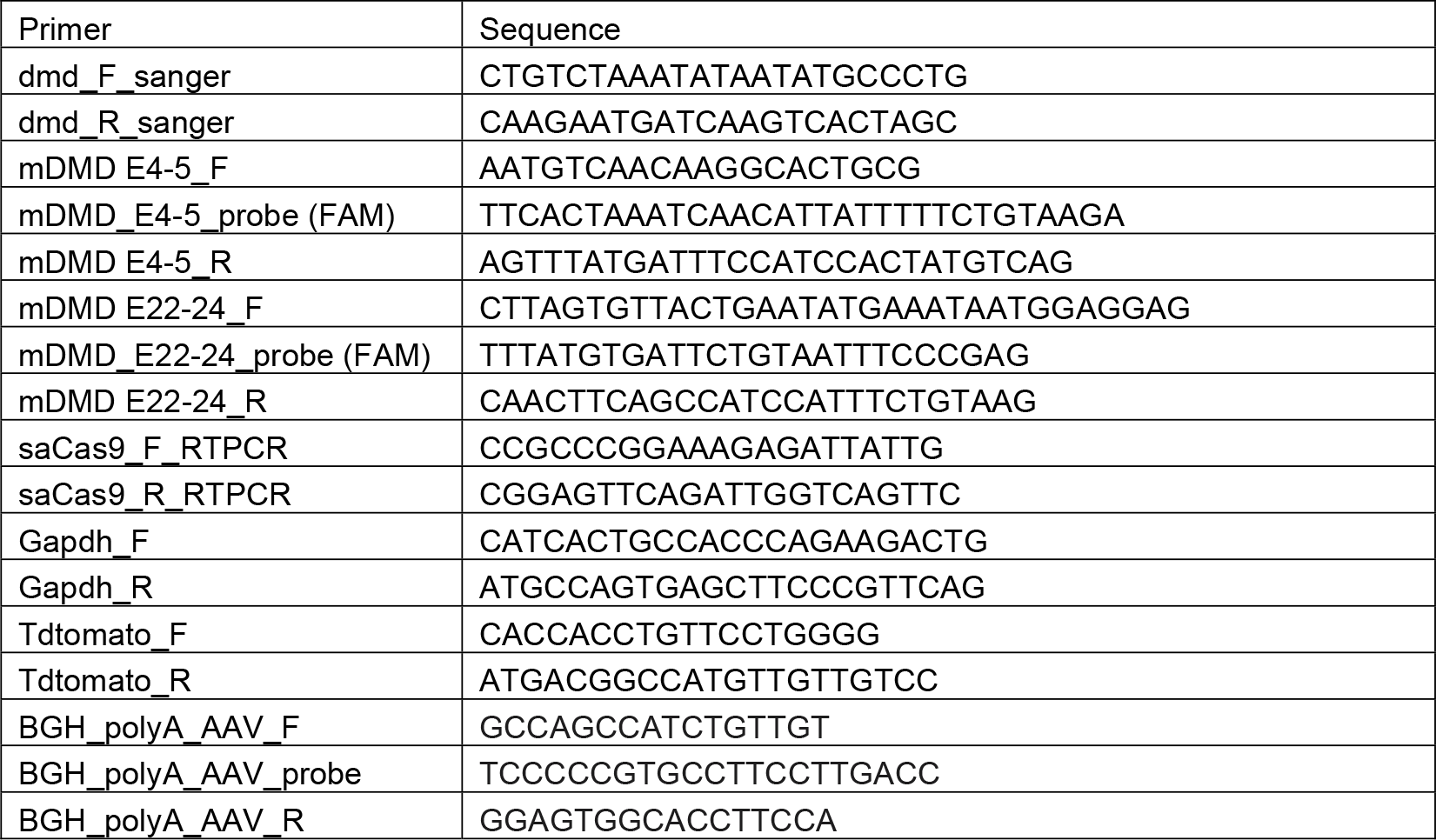
Sanger sequencing primers and qPCR primers and probes. Table listing sequences of all primers and probes used for sanger sequencing, qPCR, and RT-qPCR experiments.

**Figure S1.**
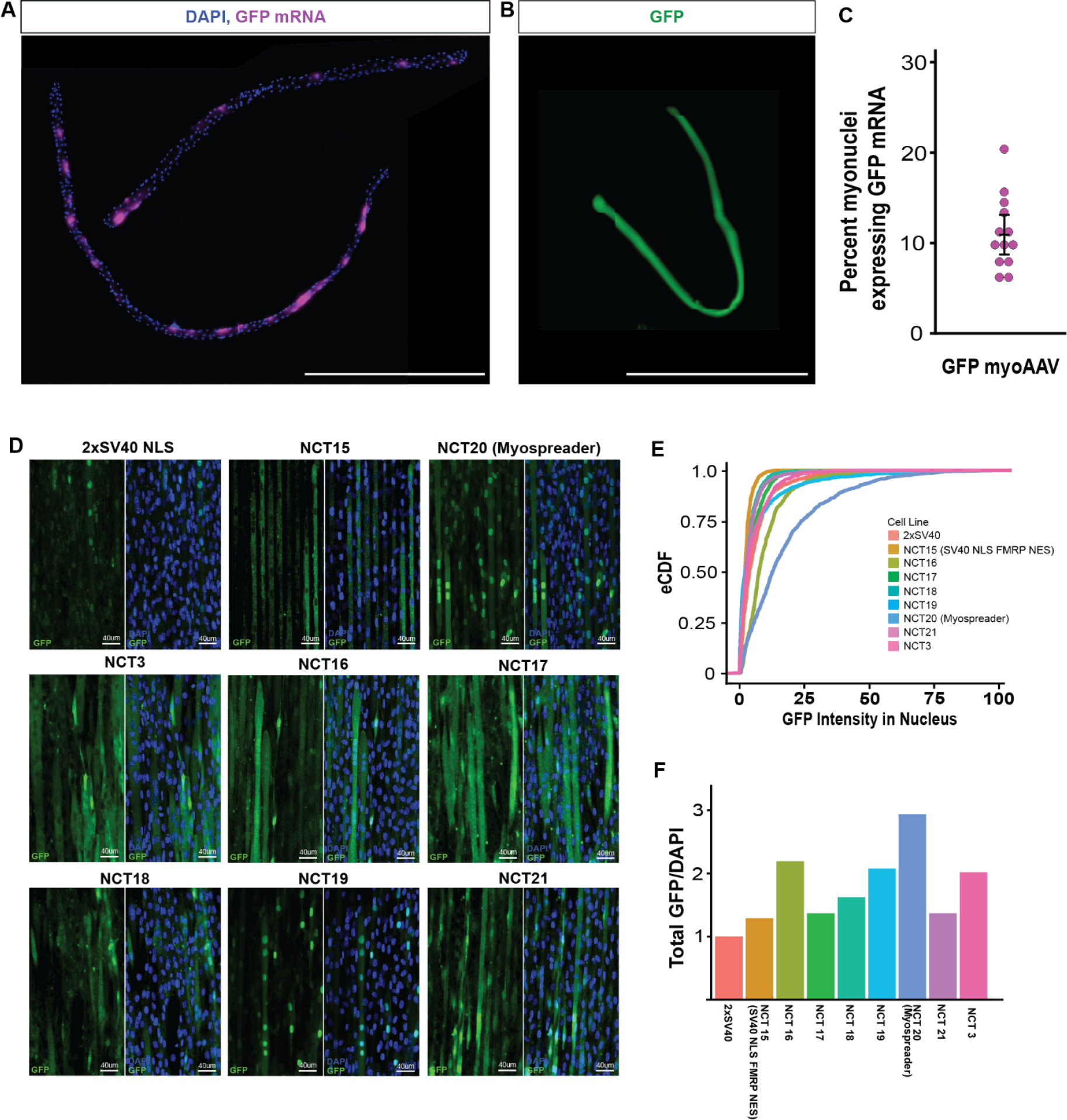
NLS/NES combinations affect GFP localization in mixed nuclei C2C12 myotubes. Related to Figure 1. (**A**) HCR-FISH to detect GFP mRNA (magenta) and DAPI (blue) in representative EDL myofibers isolated from mouse systemically treated with emyoAAV GFP. Scale bar: 1mm. (**B**) Native GFP (green) in representative EDL myofiber isolated from mouse systemically treated with emyoAAV GFP. Scale bar: 1mm. (**C**) Quantification of percentage of myonuclei in EDL myofibers expressing GFP mRNA. (**D**) Representative image of GFP fluorescence (green) in mixed nuclei C2C12 myotubes expressing additional NLS/NES combinations. Scale bar: 40µm. (**E**) Cumulative distribution functions of the nuclear GFP signal of all NLS/NES combinations tested in mixed nuclei C2C12 myotubes. (**F**) Total GFP signal divided by total DAPI signal in mixed nuclei C2C12 myotubes. Normalized to 2xSV40.

**Figure S2.**
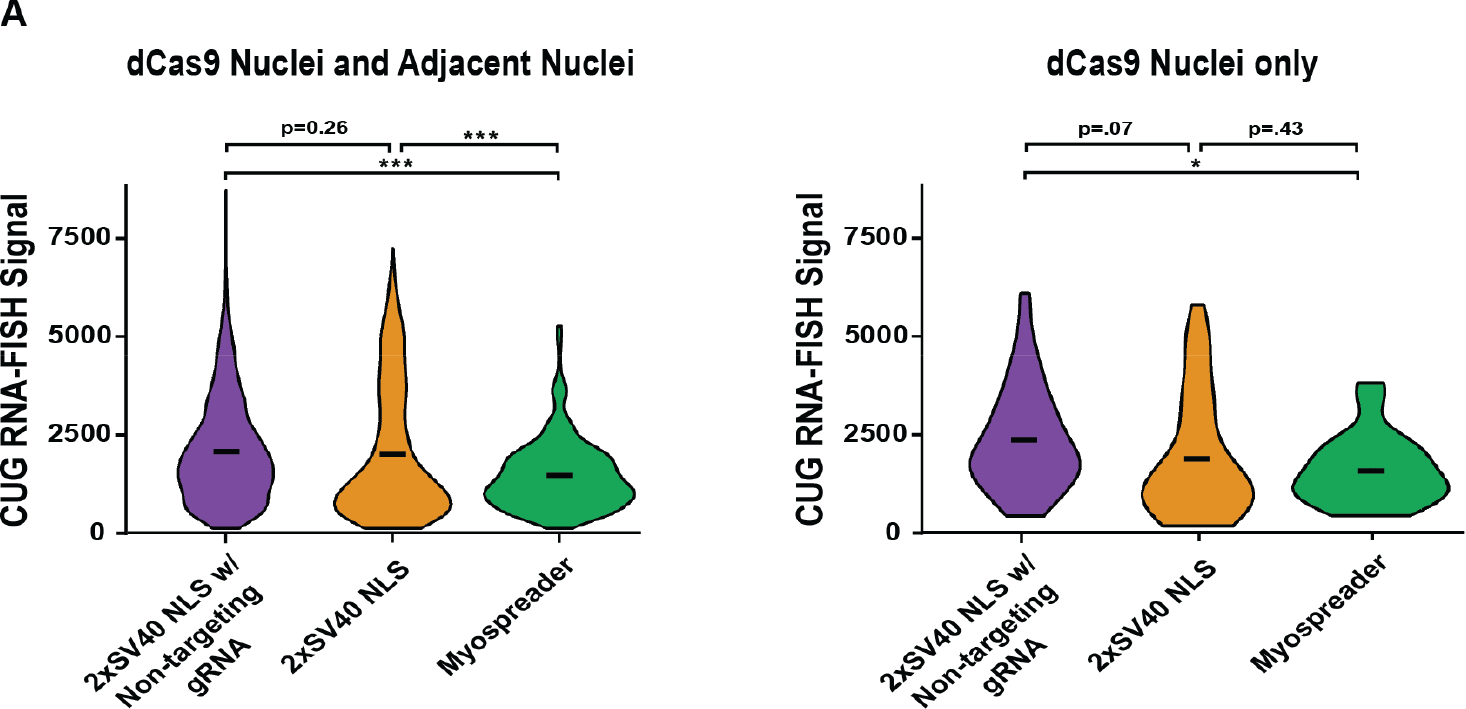
Myospreader dCas9 reduced mean CUG RNA foci signal around AAV transduced myonuclei. Related to Figure 2. (**A**) Distribution of CUG RNA signal in dCas9-expressing and proximal myonuclei compared to only dCas9-expressing myonuclei in myofibers isolated from mice treated with 2xSV40 NLS dCas9 with non-targeting gRNA AAV or 2xSV40 NLS dCas9 with CTG-targeting gRNA AAV or Myospreader dCas9 with CTG-targeting gRNA AAV. Means indicated by horizontal dashes. Significance by Student’s T-test, no correction for multiple comparisons.

**Figure S3.**
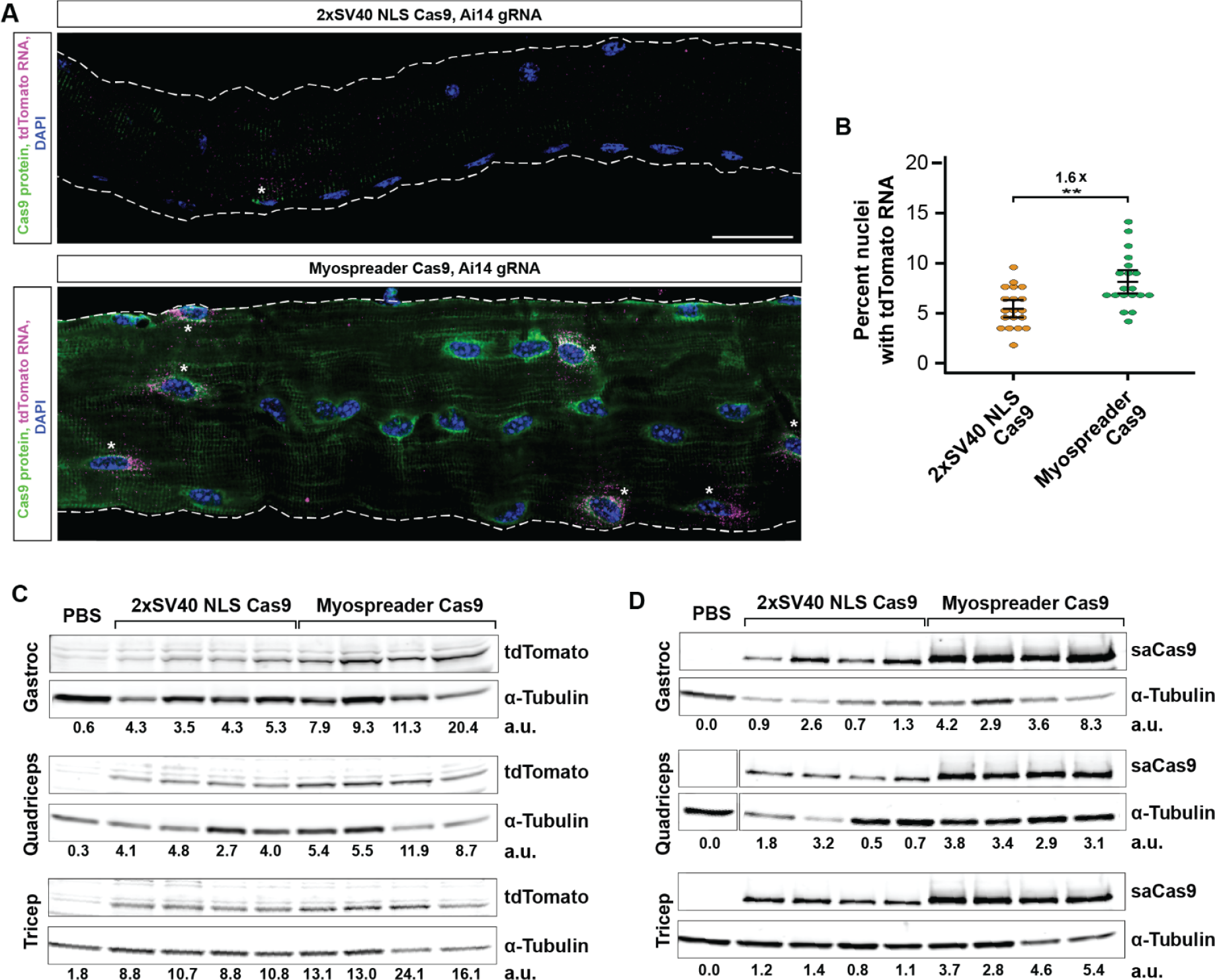
Myospreader improves DNA editing efficiency and Cas9 localization in CRISPR reporter mouse. Related to Figure 3. (**A**) HCR-FISH/IF to detect tdTomato mRNA (magenta) and Cas9 protein (green) in representative TA myofibers isolated from treated AI14 mice. tdTomato expressing nuclei indicated by asterisks. Scale bar: 30µm. (**B**) Quantitation of the percent of myonuclei expressing tdTomato mRNA detected by HCR-FISH in TA myofibers isolated from treated AI14 mice. Significance by Student’s t-test. (**C**) Western blots detecting tdTomato and α-Tubulin in muscles of treated AI14 mice. Relative signal intensity determined by densitometry is as the bottom. A.U.: Arbitraty Unit, normalized to α-Tubulin. (**D**) Western blots detecting Cas9 and α-Tubulin in muscles of treated AI14 mice. Relative signal intensity determined by densitometry is as the bottom. A.U.: Arbitraty Unit, normalized to α-Tubulin.

**Figure S4.**
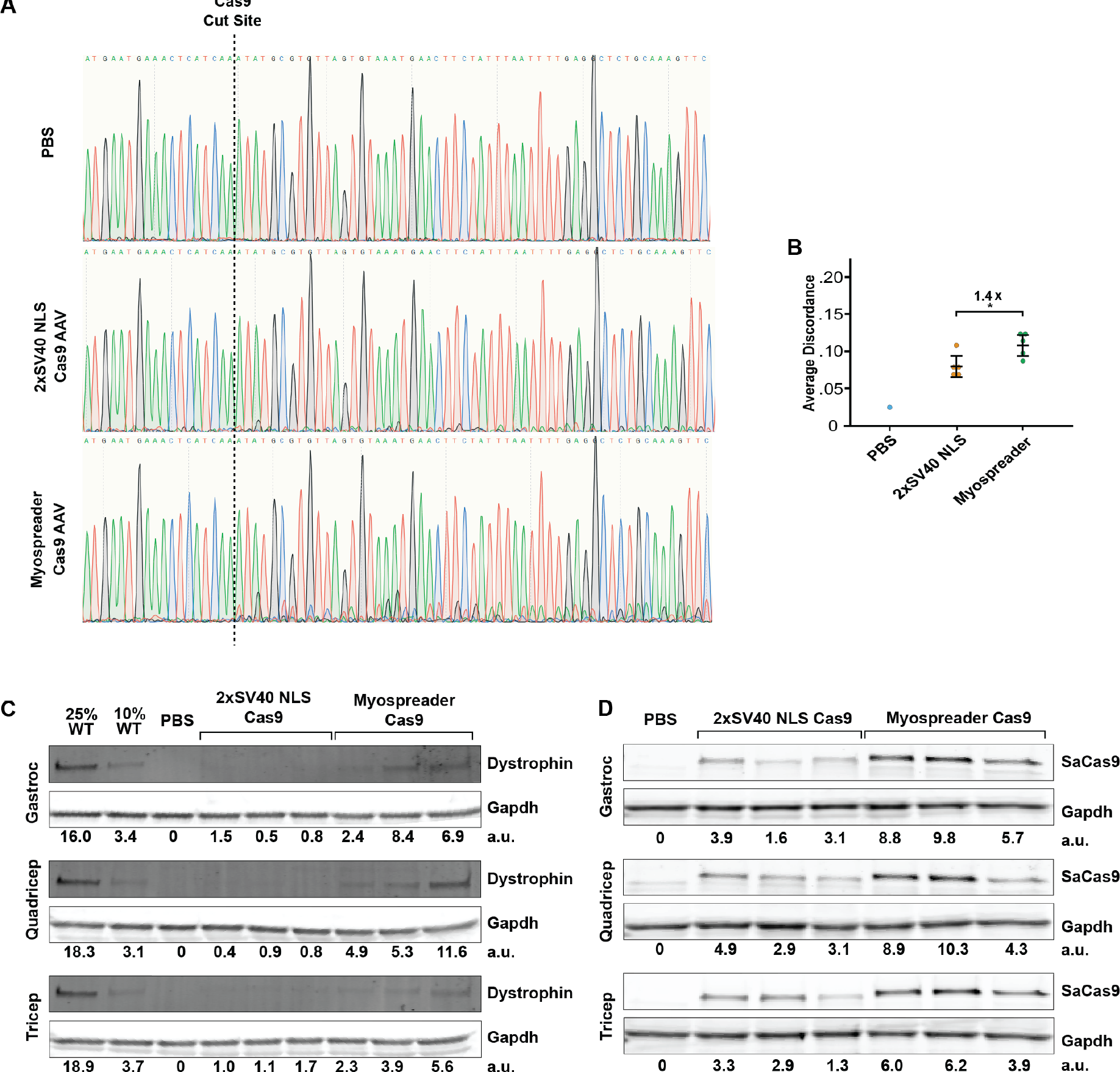
Myospreader Improves DNA editing efficiency and restores dystrophin expression in Duchenne muscular dystrophy mouse model. Related to Figure 4. (**A**) Sanger sequencing chromatogram of gastrocnemius muscle DNA of mdx mice systemically treated with PBS or 2xSV40 NLS Cas9 AAV or Myospreader Cas9 AAV. Cas9 edited DNA is observed as smaller, competing peaks after the Cas9 cut site. (**B**) Sanger sequencing average discordance of 100bp after 5’ intron gRNA target in DNA amplified from treated mdx mouse gastrocnemius gDNA. Significance by Student’s t-test. (**C**) Western blots detecting Dystrophin and GAPDH in muscles of treated mdx mice. Relative signal intensity determined by densitometry is at the bottom. A.U.: Arbitraty Unit, normalized to GAPDH. (**D**)Western blots detecting Cas9 and GAPDH in muscles of treated mdx mice. Relative signal intensity determined by densitometry is at the bottom. A.U.: Arbitraty Unit, normalized to GAPDH.

**Figure S5.**
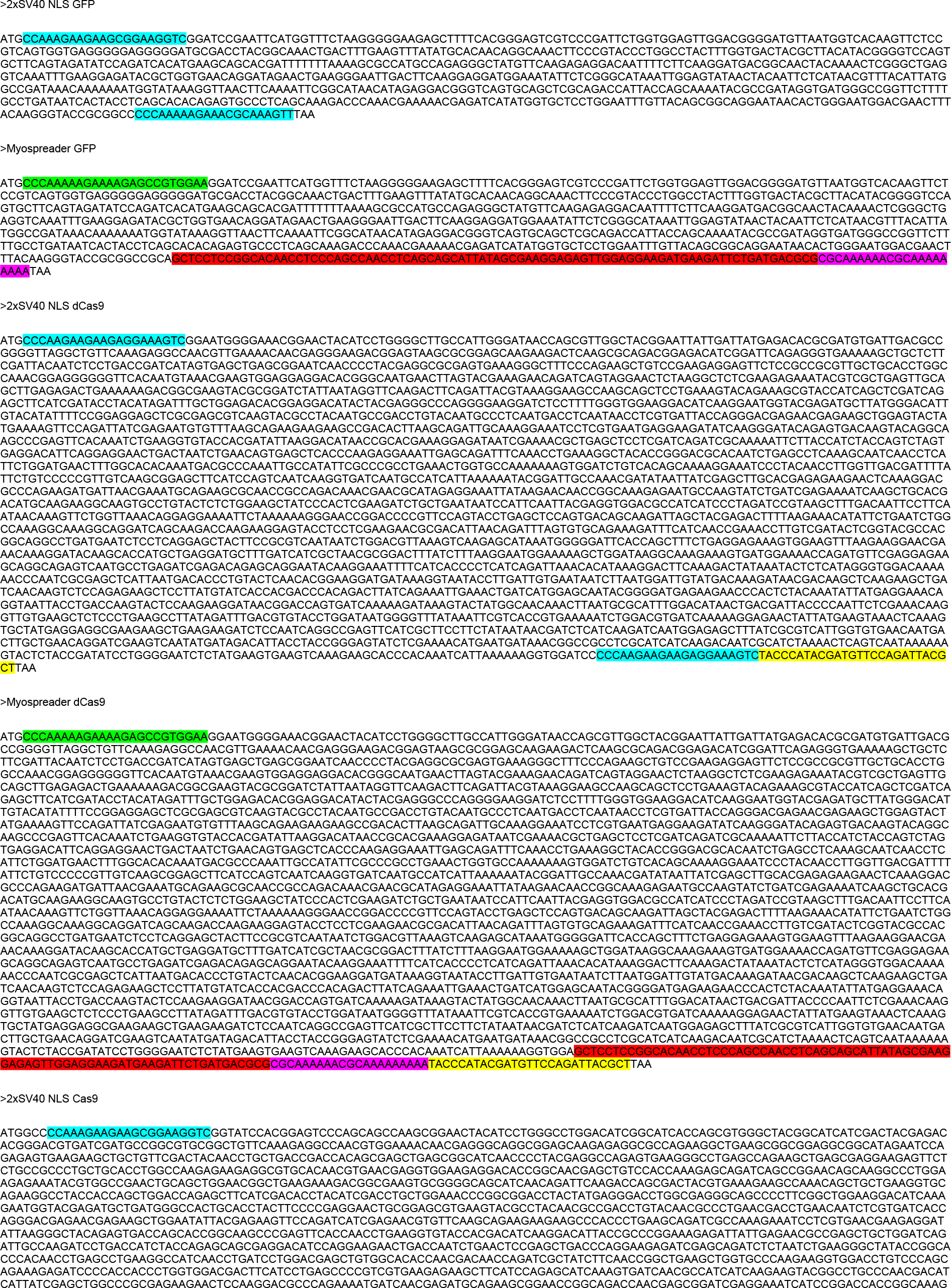
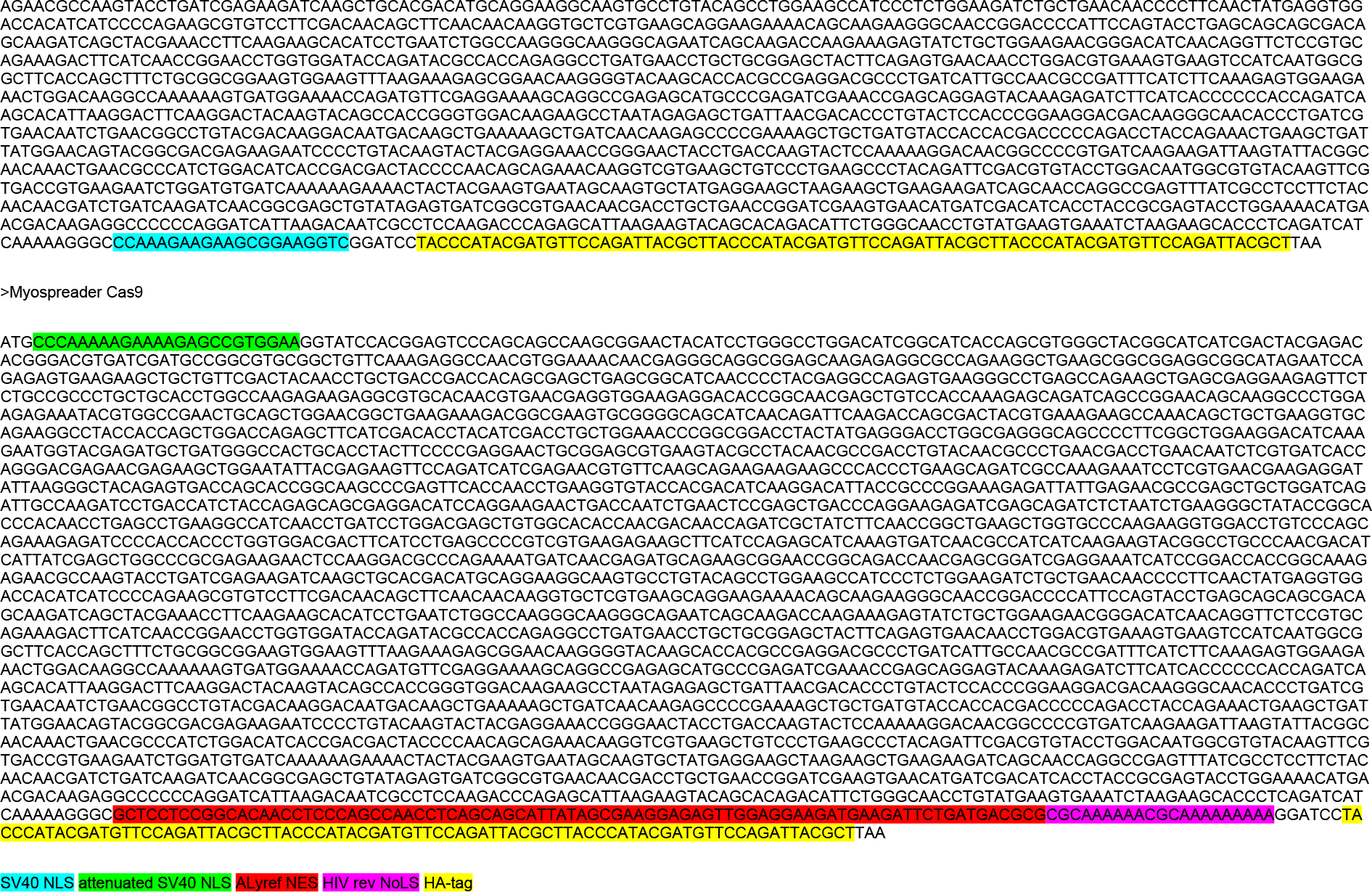
Coding sequences of AAV delivered GFP, dCas9, and Cas9. Coding sequences of AAV delivered GFP, dCas9, and Cas9 used in *in-vivo* experiments. Localization elements and HA-tags are highlighted.

